# Functional Specialization of Ca²⁺-Binding Motifs in Human MICU1

**DOI:** 10.1101/2025.06.25.661581

**Authors:** Leandro Matías Sommese, Nicolás Palopoli, Maria Silvina Fornasari, Gustavo Parisi, Toni Gabaldón, Ana Julia Velez Rueda

**Affiliations:** Structural Bioinformatics Group, Departamento de Ciencia y Tecnología, Universidad Nacional de Quilmes - Consejo Nacional de Investigaciones Científicas y Técnicas (CONICET), Buenos Aires, Argentina; Barcelona Supercomputing Centre (BSC-CNS). Plaça Eusebi Güell, 1-3. 08034 Barcelona, Spain; Institute for Research in Biomedicine (IRB Barcelona), The Barcelona Institute of Science and Technology, Baldiri Reixac, 10, 08028 Barcelona, Spain; Catalan Institution for Research and Advanced Studies (ICREA), Barcelona, Spain; CIBER de Enfermedades Infecciosas, Instituto de Salud Carlos III, Madrid, Spain

**Author notes:** Both authors contributed equally to this work (Leandro Matías Sommese, Ana Julia Velez Rueda). Corresponding author(s) (Ana Julia Velez Rueda).

**Keywords:** Ca²⁺ Uniporter, MICU1, EF-hand motif’s structural and evolutionary constraints, Conformational Diversity, Structural Dynamics

## Abstract

The mitochondrial Ca²⁺ uniporter (MCU) channel is essential for energy production, cytosolic Ca²⁺ signalling, and regulation of cell death. Its activity is regulated by the core proteins MICU1 and MICU2, which respond to intracellular Ca²⁺ levels. In cardiomyocytes, MICU1 inhibits mtMCU activity at basal Ca²⁺, with Ca²⁺-binding relieving this inhibition via a conformational change. However, the precise molecular basis for this dual regulation is unclear. While twelve MICU1 structures exist, each is approximately 30% of their structure missing, omitting key flexible regions and limits the understanding of the Ca²⁺-sensing mechanism. Here, we provide structural and computational evidence to address this gap. Using structural modelling, molecular dynamics simulations, and large-scale sequence analysis, we investigate MICU1’s Ca²⁺ binding sites from both conformational and evolutionary perspectives.

Simulations based on human MICU1 models revealed a previously uncharacterized pseudo-EF-hand (pEF-h) motif. Our findings indicate that this motif functions as an early Ca²⁺ sensor, triggering conformational transitions, including shifts in surface charge distribution and isoelectric point, that prime the canonical EF-hand sites for subsequent binding. This hierarchical activation mechanism refines MICU1’s on–off regulation of the MCU. To link this mechanism to experimental observations, we simulated a series of point and double mutants targeting the pEF-h, EF-h1, and EF-h2 sites. Our simulations demonstrate that double mutants disrupt Ca²⁺ binding not only within the mutated site but also reduce the occupation of the other sites, reaffirming the cooperative nature of Ca²⁺ sensing in MICU1.

The biological relevance of the EF-hand motifs would be supported by its evolutionary conservation. Therefore, we analysed the evolutionary shaping of MICU1 EF-hand motifs across major eukaryotic lineages using clustering analysis and found strong lineage-specific segregation: canonical DXN/DXD-type motifs predominated in EF-h1 and EF-h2 in plants and protists, while non-canonical EXE(X)₃DEG(X)₇E motifs were exclusive to Opisthokonts, coinciding with the emergence of the auxiliary subunit EMRE. This pattern suggests that high-affinity Ca²⁺ binding evolved in parallel with increasing regulatory complexity in metazoans.

Together, these findings support previous research linking EF-hand function as sensors to specialised Ca²⁺ gatekeepers in multicellular lineages. By integrating structural and evolutionary perspectives, our study provides mechanistic insight into how MICU1 can act as a Ca²⁺-dependent molecular switch, clarifying the cooperative and threshold-setting behaviour underlying its regulatory role in mitochondrial Ca²⁺ uptake.

**Highlights:**

- The hierarchical activation mechanism refines MICU1’s on–off regulation of the MCU.
- A novel Ca²⁺-binding site in human MICU1, featuring a unique helix–loop–β-sheet structure, was identified and functionally characterized.
- This new Ca²⁺-binding site in MICU1 acts as an early sensor, triggering structural changes key to its regulatory function.
- Ca²⁺-binding motif distributions and phylogenetic constraints indicate possible functional divergence.

## Introduction

The mitochondrial uniporter is a Ca²⁺ channel complex (mtCU) with important roles in energy production, cytosolic Ca²⁺ signalling, and cell death [19]. In mammals, the uniporter complex comprises four core proteins: the pore-forming MCU, the gatekeepers MICU1 and MICU2, and the auxiliary subunit EMRE, which is essential for Ca²⁺ transport [4:2:2:4] [56]. Tight regulation of the uniporter by cytosolic Ca²⁺ levels is crucial for maintaining physiological homeostasis [13,17,49].

MCU homologues are widely distributed across all major branches of the eukaryotic tree of life. In sequenced metazoa, plants, and protozoa, homologues of both MCU and MICU1 are typically either present or absent together, with mammals representing an exception, as the two proteins physically interact and show high co-expression [7]. Previous studies have proposed that the co-occurrence of MCU and MICU1 may have been evolutionarily advantageous in eukaryotes [53,58]. MICU1 appears to promote organismal fitness by enhancing the selectivity of the mitochondrial Ca²⁺ uniporter, thereby preventing toxic manganese accumulation and oxidative stress, while simultaneously supporting efficient Ca²⁺-dependent ATP production in tissues with high energy demands. In mammals, MICU1 is also physically associated and co-expressed with MICU2, whose activity and physical association with the pore depend on the presence of MICU1.

The activity of the entire channel is known to be regulated by Ca²⁺ concentration through the concerted action of MICU1 and MICU2. In addition, it has been demonstrated that cells lacking MICU1 or MICU2 lose the normal threshold for Ca²⁺ intake [26]. In resting cardiomyocytes, Ca^2+^ is not bound to mitochondrial MICU1 and its binding inhibits MCU. Ca²⁺ release from the endoplasmic reticulum leads to an increase in cytosolic Ca²⁺ concentration. Upon cardiomyocyte contraction, MICU1 binds Ca²⁺ thereby activating the MCU channel, which triggers in massive Ca²⁺ uptake by mitochondria. The resulting decrease in cytosolic Ca²⁺ concentration below the activation threshold causes Ca²⁺ dissociation from MICU1, inhibiting the MCU channel [41].

MICU1 confers cooperative activation of the mitochondrial Ca²⁺ uniporter once cytosolic Ca²⁺ surpasses a specific threshold, ensuring appropriate mitochondrial Ca²⁺ uptake in response to cellular signals [10]. Csordás and co-authors compared mitochondrial Ca²⁺ uptake differences between wild type and MICU1-deficient cells, and demonstrated that its two functional Ca²⁺-binding EF-hand motifs (EF-h) in the protein are important in the transition between fully closed and open states of MCU. Even though the authors suggested that MICU1 conformational changes may mediate the regulation of multiple specific functions required in different cell contexts, the underlying structural mechanisms of this switch mechanism remain unknown. The mtCU’s Ca²⁺-dependent regulation hinges on dynamic interactions among MCU, MICU1, MICU2, and EMRE. In low intracellular Ca²⁺ concentration (<1 μM), MICU1’s conserved K/R ring (residues K126, R128, R129, R130, R132) binds MCU’s acidic pore entrance (D261, E264), blocking Ca^2+^ flux, while its N-terminal polybasic domain (residues 99–110) and C-terminal helix (residues 440–443, including R440/R442) stabilize the MCU-MICU1 interface. MICU1 further anchors to EMRE via electrostatic interactions between its polybasic region and EMRE’s acidic C-terminal tail (D68/E72). Both MICUs form a Ca²⁺-sensing heterodimer through critical EF-h interactions (between MICU1 D231/D235 and MICU2 R352). When Ca²⁺ concentration rises, cooperative binding to the MICU1-MICU2 heterodimer induces a 30° rotation of MICU1’s EF1 helix (α6), displacing pore-contacting residues (R259/R261/R263) and reorienting the K/R ring (K126/R129), thereby unblocking the pore. Throughout this gating cycle, EMRE maintains MICU1 proximity to MCU, ensuring rapid channel re-closure upon Ca²⁺ decrease [16].

While MICU1 is widely recognised as the primary gatekeeper of the mitochondrial Ca²⁺ uniporter, there is ongoing debate regarding the specific contributions of MICU1 and MICU2 to its switch-like behaviour. Several studies indicate that MICU1 alone is sufficient to establish the basal Ca²⁺ threshold, preventing uniporter opening at resting cytosolic Ca²⁺ levels [11,13]. In contrast, other reports suggest that MICU2 is essential for the cooperative activation observed at higher Ca²⁺ concentrations, modulating the steepness and sensitivity of the uniporter’s response [9,10]. Structural analyses have demonstrated that MICU1 physically blocks the MCU pore under low Ca²⁺ conditions [2,14], whereas the conformational contributions of MICU2 during activation remain less clearly defined. From an evolutionary perspective, MICU1 is broadly conserved across eukaryotes and likely represents the ancestral gatekeeper, while MICU2 appears only in metazoans, suggesting that it provides additional regulatory refinement to the core MICU1-mediated threshold control [15,16]. Addressing the remaining questions about MICU1’s gating mechanism requires combining evolutionary analysis with structural and computational approaches. Comparative studies of MICU1 homologues across eukaryotes can identify conserved residues and motifs critical for its Ca²⁺-dependent on–off switching, while molecular dynamics simulations can provide mechanistic insights into the conformational changes triggered by Ca²⁺-binding. Together, these approaches can clarify the structural basis of its threshold regulation and cooperative activation.

MtCU has been associated with different diseases [2,15,51], highlighting its clinical relevance in human health. Particularly, MICU1 dysfunction leads to severe pathological conditions due to impaired mitochondrial Ca²⁺ homeostasis [6]. In Parkinson’s disease, given that MICU1 is crucial for neuronal survival by regulating Ca²⁺ efflux from mitochondria via the Na⁺/Ca²⁺ exchanger, its dysfunction leads to mitochondrial Ca²⁺ overload and increased vulnerability to cell death [15]. Studies in pancreatic β-cells have demonstrated that both MICU1 and MCU are essential for proper insulin secretion, as their knockdown significantly reduces glucose-stimulated insulin secretion by affecting mitochondrial Ca²⁺ uptake and ATP production [2]. Moreover, in diabetic cardiomyopathy, MICU1 deficiency in cardiac microvascular endothelial cells leads to excessive mitochondrial Ca²⁺ uptake and homeostasis imbalance, causing nitrification stress-induced endothelial damage and inflammation that disrupts the myocardial microvascular endothelial barrier function [51].

Loss-of-function mutations in MICU1 result in a complex disease related phenotype characterized by proximal muscle weakness, learning difficulties, and extrapyramidal motor disturbances [8,31]. To date, 13 pathogenic MICU1 variants have been identified, with mutations occurring through different mechanisms including splice site mutations, nonsense variants, missense variants, and deletions [8,29]. However, the structural basis underlying the functional impact of many of these mutations is still unclear. In particular, the precise conformational transitions that enable MICU1 to switch between inhibitory and permissive states in response to Ca²⁺ remain incompletely characterized, especially in the context of its interaction with other components of the MCU complex [26,41]. Additionally, the contribution of non-canonical motifs to MICU1’s sensing and gating behavior has not been fully explored in vivo. Moreover, how tissue-specific expression and post-translational modifications modulate MICU1 activity across different physiological and pathological conditions is still largely unknown [10,30]. This is especially relevant given experimental evidence showing that even single mutations in non-EF-hand regions, such as R185 deletion, can produce subtle effects, while combined mutations generate profoundly disruptive phenotypes (e.g. D421A+E432K or D231A+E242K mutants), revealing the cooperative nature of MICU1 Ca²⁺ sensing [27,42,59]. Addressing these gaps is essential to fully decipher MICU1’s role in mitochondrial signaling and to develop targeted interventions for MICU1-related disorders.

In this study, we identified and characterized a previously unrecognized pseudo-EF-hand (pEF-h) motif in MICU1 that may be involved in gatekeeping regulation. Through computational modeling and molecular dynamics simulations, we describe the conformational changes induced by Ca²⁺-binding and reveal a stepwise activation mechanism. Our results indicate that the pEF-h acts as an early Ca²⁺ sensor, triggering structural rearrangements that facilitate subsequent Ca²⁺-binding at the canonical EF-hand sites. These conformational shifts affect charge distribution, protein flexibility, and intramolecular interactions, ultimately modulating MICU1’s gating function over mitochondrial Ca²⁺ uptake. Furthermore, we observed a progressive decrease in Gibbs free energy (ΔG) upon Ca²⁺ binding—first at the pEF-h, then at the canonical EF-hand sites—suggesting a cooperative mechanism in which initial binding events stabilize the protein fold and enhance overall Ca²⁺ affinity.

Our evolutionary analysis of MICU1 homologs showed how the Ca²⁺ binding sites shaped the functional needs of each species. The pEF-h and other Ca²⁺ binding sites across major eukaryotic lineages underwent lineage-specific segregation: non-canonical DXN/DXD motifs predominated in basal groups, whereas canonical EXE motifs were enriched in Opisthokonta, coinciding with the emergence of EMRE. By identifying this novel regulatory element, our findings enhance the understanding of MICU1’s structural dynamics and functional regulation, paving the way for future studies on its role in mitochondrial Ca²⁺ homeostasis and its involvement in conditions such as diabetic cardiomyopathy and neurodegenerative diseases.

## Results

### Human MICU1 functional and structural characterization

The regulatory subunit MICU1 is a large protein, comprising 476 amino acids and weighing approximately 54 kDa. The region between residues 1 and 110 undergoes order–disorder transitions. This region forms a compositionally biased sequence subdomain, with basic and acidic residues at the MCU–MICU1 interface [21]. As all MICU1 three-dimensional structures available in the Protein Data Bank (PDB) are incomplete [5], we modelled the 30% missing portion of MICU1 using AlphaFold2 [25] followed by molecular dynamics minimisation. To address persistent knowledge gaps regarding the conformational transitions that enable MICU1’s regulatory function, we examined the conformational changes upon Ca²⁺ binding. Inspection of the human MICU1 model revealed three Ca²⁺-binding sites: two previously described canonical EF-hands (EF-hand 1 and EF-hand 2), displaying the conserved DXNXDG(X)5E/DXDXNG(X)5E linear motif and the typical α-loop-α structure, and a third, previously unknown site. This novel binding site is present in both the fully unbound and fully Ca²⁺-bound forms and is located within a region between Glu184 and Glu200 that is absent from the available PDB structures. We characterized this region as a pseudo-EF-Hand motif (pEF-h), as its structural configuration deviates from the canonical EF-h motif [25] —the canonical linear motif DXDXNG(X)5E with an **α**-loop-**α** structure—having instead the linear motif EXE(X)3DEG(X)7E with an **α**-loop-**β** structure (see Figure 1). Despite this structural divergence, the site facilitates Ca²⁺ coordination and appears to play a crucial role in the protein’s Ca²⁺ handling, as well as the structural and functional changes driven by Ca²⁺-binding, as suggested by the results presented below.

**Figure 1.**
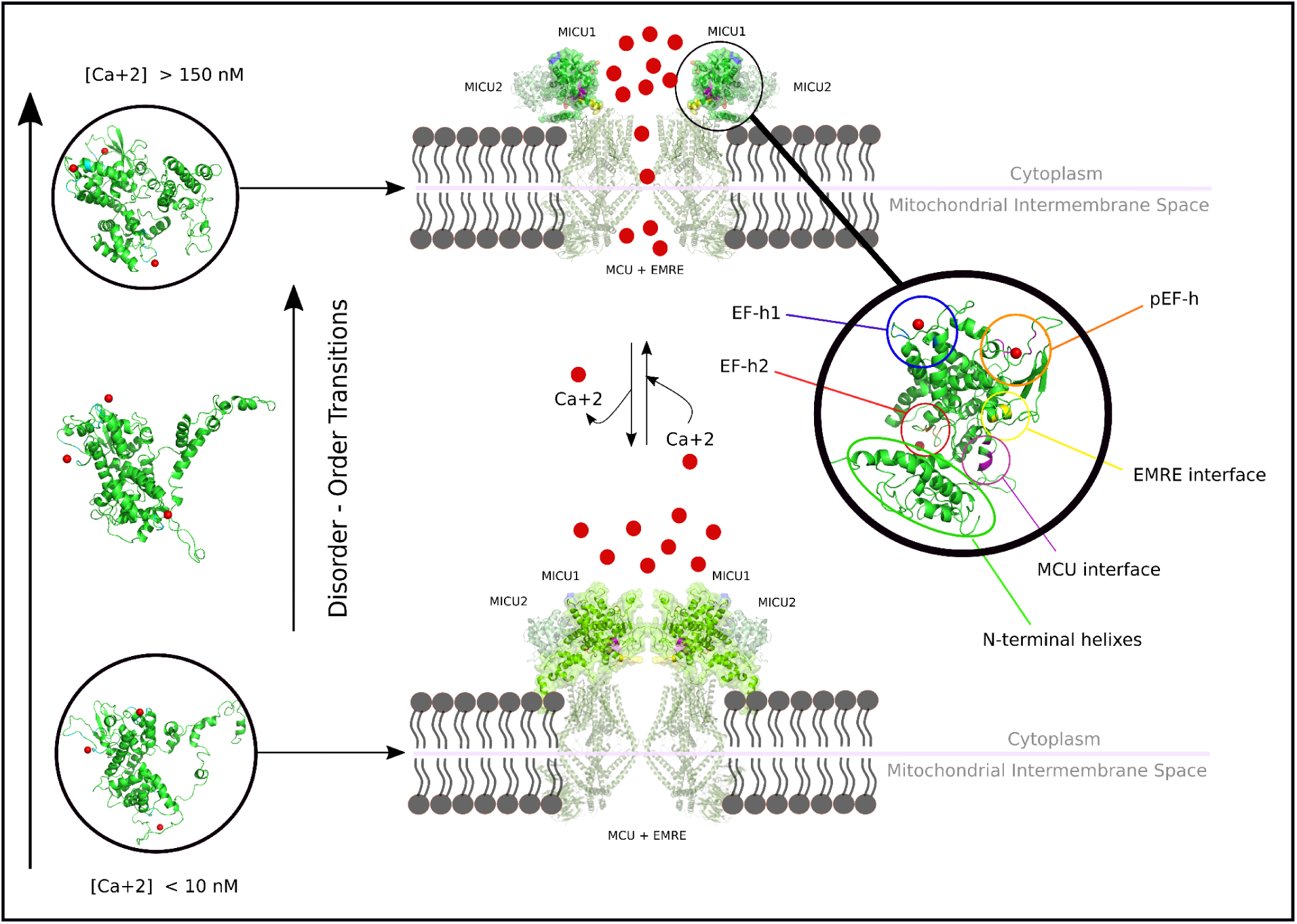
Schematic representation of the MICU1 mediated Ca^2+^-sensing switch in the mitochondrial Ca^2+^ uniporter (mtCU) complex. The diagram illustrates the key conformational and functional states of the MICU1-MICU2 regulatory dimer relative to the MCU-EMRE pore. Ca^2+^ ions are shown as red spheres. At the top, at high cytosolic Ca²⁺ concentrations ([Ca²⁺] > 150 nM), Ca²⁺ binding to MICU1 triggers disorder-to-order transitions, particularly in the N-terminal helices. This results in a more compact MICU1 conformation and causes the MICU1-MICU2 complex to move away, exposing the EMRE and MCU interfaces. This allows EMRE-MCU engagement and pore opening for Ca²⁺ uptake into the mitochondrial matrix. At the bottom, at low, resting Ca²⁺ concentrations ([Ca²⁺] < 10 nM), MICU1 exhibits disordered regions (wavy lines) in its N-terminus and does not achieve full structural stabilization due to incomplete Ca²⁺ occupancy at the EF-hand sites. In this state, the MICU1-MICU2 dimer maintains a physical blockade over the EMRE interaction site on MCU, preventing pore opening and keeping the channel closed. Key Ca²⁺-binding sites are highlighted in MICU1 tridimensional structure: blue indicates the canonical EF-hand 1 (from D231 to E242), while red marks EF-hand 2 (from D421 to E432), located near the MICU2 binding site (from F383 to Q398). The purple region highlights the MCU interaction interface (from F106 to Y121), and the yellow region denotes the EMRE interaction site (from K126 to R129).

The Janus analysis [32,40], aimed at identifying metamorphic proteins that exhibit fold-switching depending on environmental conditions, revealed a sequence with context-dependent conformational flexibility, expected to collapse or expand based on environmental conditions [32]. Given the hypothesis that MICU1 functions as an on-off switch depending on Ca²⁺ concentration [60], we computationally simulated the titration of MICU1 by performing molecular dynamics in solvation solutions containing 0, 10, 50, and 150 mM Ca²⁺. We observed structural differences between the unbound and Ca²⁺-bound states of MICU1. within the first 30 nanoseconds of the dynamics. The interaction with Ca²⁺ at 150 mM induced significant structural changes (RMSD = 13.17 Å) (Figure 1), reducing its theoretical isoelectric point from 10.03 to 2.93 and changing the protein’s net charge from +1 to -1 under the same pH and ionic strength conditions. Additionally, Ca²⁺ binding decreased the volume, length, and hydrophobicity of all MICU1 cavities by 14.28%, 27.5%, and 41.7%, respectively. After the initial transition, the Ca²⁺-bound structure stabilized, and no further major conformational changes were observed for the remainder of the simulation, with RMSD plateauing around 17,7 Å.

One of the most interesting observations was the order–disorder transition in the N-terminal region, guided by Ca²⁺-binding (Figure 1). This negatively charged polyampholyte region forms three alpha helices capable of coordinating multiple Ca²⁺ ions through electrostatic interactions with non-conserved negative residues. The distances between the EMRE-binding alkaline motif and the Ca²⁺-binding site EF-h2 decrease substantially. For example, the most pronounced changes are observed between residues K351 and D421, which decrease from 13.0 to 11.3 Å after binding to Ca²⁺, and between residues K340 and E432, which decrease from 19.2 to 13.9 Å (Figure 2A). The presence of Ca²⁺ promotes the reorientation of residues K340 and K341, disrupting the interactions maintained with the MICU II/EMRE site (polylysine region) through hydrogen bonding with residues D78, E79, and E432 (Figure 2C). This movement ultimately exposes interaction region II, facilitating binding between these two proteins in a new orientation. This structural change, triggered by Ca²⁺-binding, allows the exposure and reorientation of binding sites to MCU and EMRE (Figure 2B).

**Figure 2.**
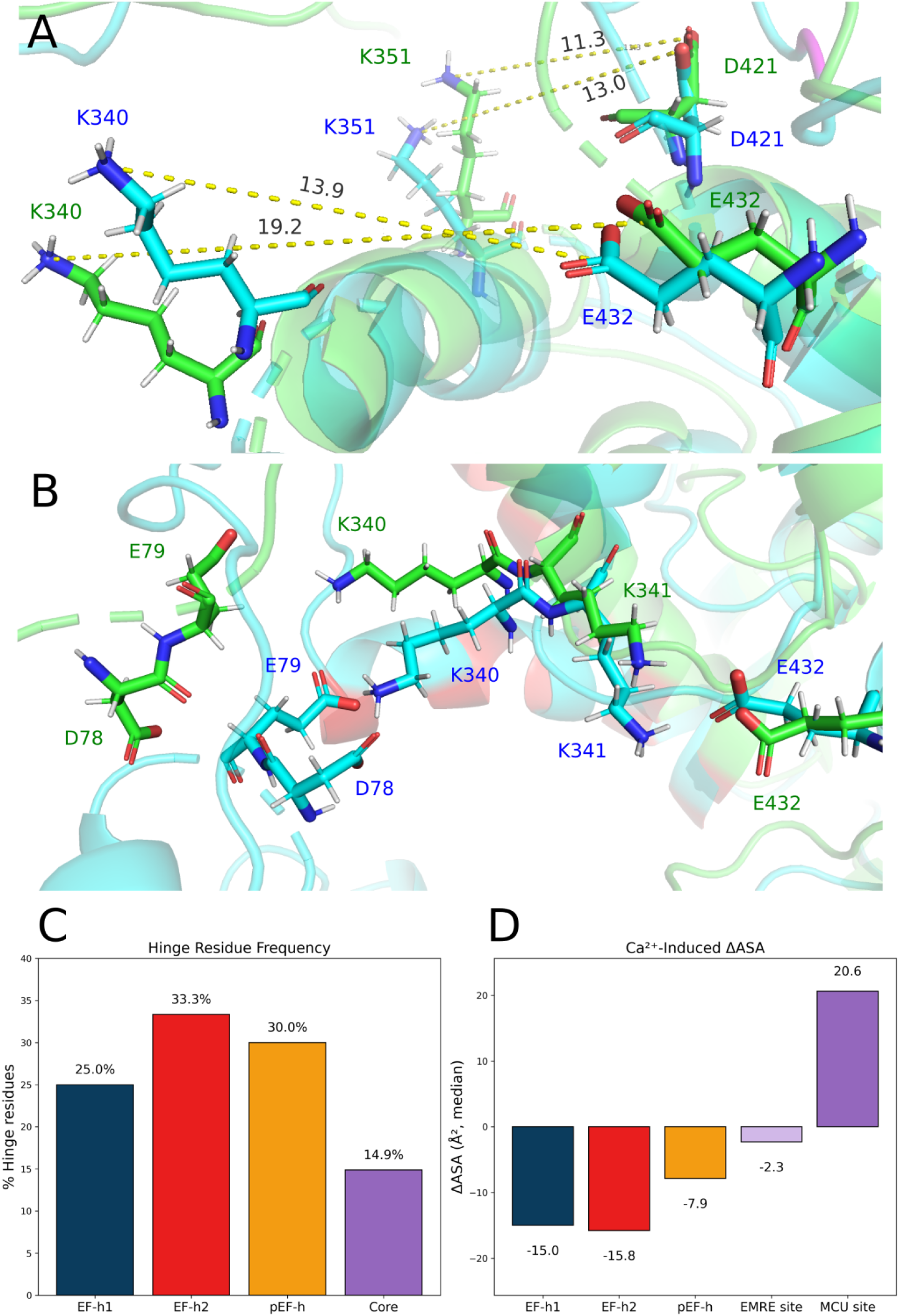
Ca^2+^-binding (cyan) induces structural rearrangements that reduce distances between the EMRE-binding alkaline motif and Ca²⁺ site II, compared to the unbound state (green); 2A) Notable reductions occur between K351–D421 (13.0→11.3 Å) and K340–E432 (19.2→13.9 Å); 2B) This reorientation disrupts interactions with MICU1/EMRE (D78, E79, E432), exposing interaction region II and enabling new MCU-EMRE binding; 2C) Hinge residue frequency (%) per region, calculated from torsion angles: regions with the largest dihedral changes (EF-h1, EF-h2, and pEF-h motifs) are enriched in hinge residues, while the core shows moderate flexibility. 2D) ΔASA Bound – Unbound median (Å²) for the same regions, showing the effect of Ca²⁺ binding on solvent accessibility. EF-hand and pEF-hand regions decrease in ASA (conformational collapse), whereas the MCU interface increases, indicating exposure of this interaction site. Colors in both 2C and 2D correspond to: blue for EF-h1, red for EF-h2, orange for pEF-hand, and purple for the protein core (including EMRE and MCU sites).

Torsion angle analysis showed that Ca²⁺-induced conformational changes in MICU1 are not uniformly distributed along the protein but are mediated by discrete backbone rotation sites that act as mechanical hinges. Hinge residues, defined as those in the upper 25th percentile of total backbone dihedral variation between unbound and Ca²⁺-bound states, were significantly enriched in the EF-hands and pEF-hand regions. Among these, the pEF-hand exhibited the largest mean dihedral change (51.7°) and one of the highest fractions of hinge residues (30%), identifying it as the dominant conformational transducer. In contrast, the structural core showed smaller dihedral changes and a lower proportion of hinge residues, consistent with its role as a relatively rigid scaffold (Figure 1C). These results support a hinge-driven allosteric mechanism in which Ca²⁺ binding induces localized backbone rotations within the EF-hand and pEF-hand motifs that are transmitted through the protein framework to modulate interaction surfaces, rather than those surfaces undergoing large intrinsic rearrangements themselves. Consistently, analysis of solvent accessibility (ASA) revealed that Ca²⁺ binding reduces the median ASA of the EF-hand and pEF-hand regions, reflecting their conformational collapse after Ca²⁺ binding, while increasing the ASA at the MCU interface, supporting the exposure of this interaction site (Figure 1D). In contrast, the EMRE site shows minimal ASA changes, likely because it is already solvent-exposed and structurally flexible in both unbound and Ca²⁺-bound states, making ΔASA a less sensitive indicator for this region.

Interestingly, we observed a reduction in intermolecular distance between the coordinating atoms of EF-hand sites 1 and 2, with a substantial decrease of 53.4% upon transition from the unbound to the fully Ca²⁺-bound forms (see Table 1). Similarly, the pEF-h site also undergoes a significant reduction in intermolecular distance upon Ca²⁺-binding, from 11.3 Å to 6.6 Å, highlighting its role in Ca²⁺ coordination (see Table 1). Notably, as the Ca²⁺ concentration increases from 10 mM to 150 mM, the Ca²⁺-binding sites exhibit cross-activation between the unbound and fully Ca²⁺-bound forms, along with a pKa shift of -0.55/0.33.

**Table 1.** Distances between residues within Ca^2+^-binding sites between unbound (green) and and the fully bound (cyan) conformation, indicating the spatial relationships during unbound and fully Ca²⁺-bound states. The values are measured in angstroms (Å) from the coordinating atoms of each residue and represent the proximity of specific residue pairs involved in Ca^2+^ coordination.

|  | EF-h1 |  |  | EF-h2 |  |  |  | pEF-h |
| --- | --- | --- | --- | --- | --- | --- | --- | --- |
|  | D231/<br>N233 | N233/<br>D235 | K290/<br>E242 | D423/<br>E427 | N425/<br>E432 | D421/<br>E432 | D421/<br>D423 | S138/<br>E200 |
| Unbound | 7.2 $\text{\AA}$ | 5.9 $\text{\AA}$ | 7.1 $\text{\AA}$ | 12.9 $\text{\AA}$ | 11.6 $\text{\AA}$ | 5.9 $\text{\AA}$ | 5.9 $\text{\AA}$ | 11.3 $\text{\AA}$ |
| Fully<br>$\text{Ca}^{2+}$-bound | 2.9 $\text{\AA}$ | 4.4 $\text{\AA}$ | 5.4 $\text{\AA}$ | 8.0 $\text{\AA}$ | 6.3 $\text{\AA}$ | 8.7 $\text{\AA}$ | 5.4 $\text{\AA}$ | 6.6 $\text{\AA}$ |

### Regulation of MICU1 Function and Energetics Through Ca^2+^ Binding

Furthermore, to understand the functional role of these three sites in the on-off switch transitions, we evaluated their stickiness and occupancy. For the first metric, we measured the duration each Ca^2+^ ion remained bound to a site; for the second, we assessed the proportion of residues at each site that maintained contact with Ca^2+^. Molecular dynamics analysis revealed that at high Ca^2+^ concentrations ([Ca²⁺] > 50 mM), there is permanent occupation of pEF-h and EF-h1 throughout the entire trajectory, and a late occupation of EF-hand2, which precedes the protein disorder–order transition in the N-terminal region (Figure 3). At lower Ca^2+^ concentrations ([Ca²⁺] = 10 mM), EF-h2 does not exhibit permanent Ca^2+^ occupation, although both the pEF-h and EF-h1 sites establish contacts with the ion. Notably, even though the pEF-h site binds Ca^2+^ earlier than EF-h1, the latter maintains more persistent contacts. The lack of complete binding at low Ca^2+^ concentrations does not allow full protein folding, reducing the permanence and occupancy of the sites. In summary, we observed a temporal sequence in Ca^2+^-binding: Ca^2+^ first binds to the pEF-h site, which acts as a sensitive Ca^2+^ sensor. This initial binding induces conformational changes that subsequently enable the EF-hand sites to bind Ca^2+^. These findings suggest that both pEF-h and EF-hand sites play crucial roles in the open–close switching mechanism of the channel and provide a structural mechanism for experimental observations in which Ca²⁺-binding to MICU1 EF-hands induces cooperative activation of mtCU, and contributes with temporal channel regulation [10,41].

**Figure 3.**
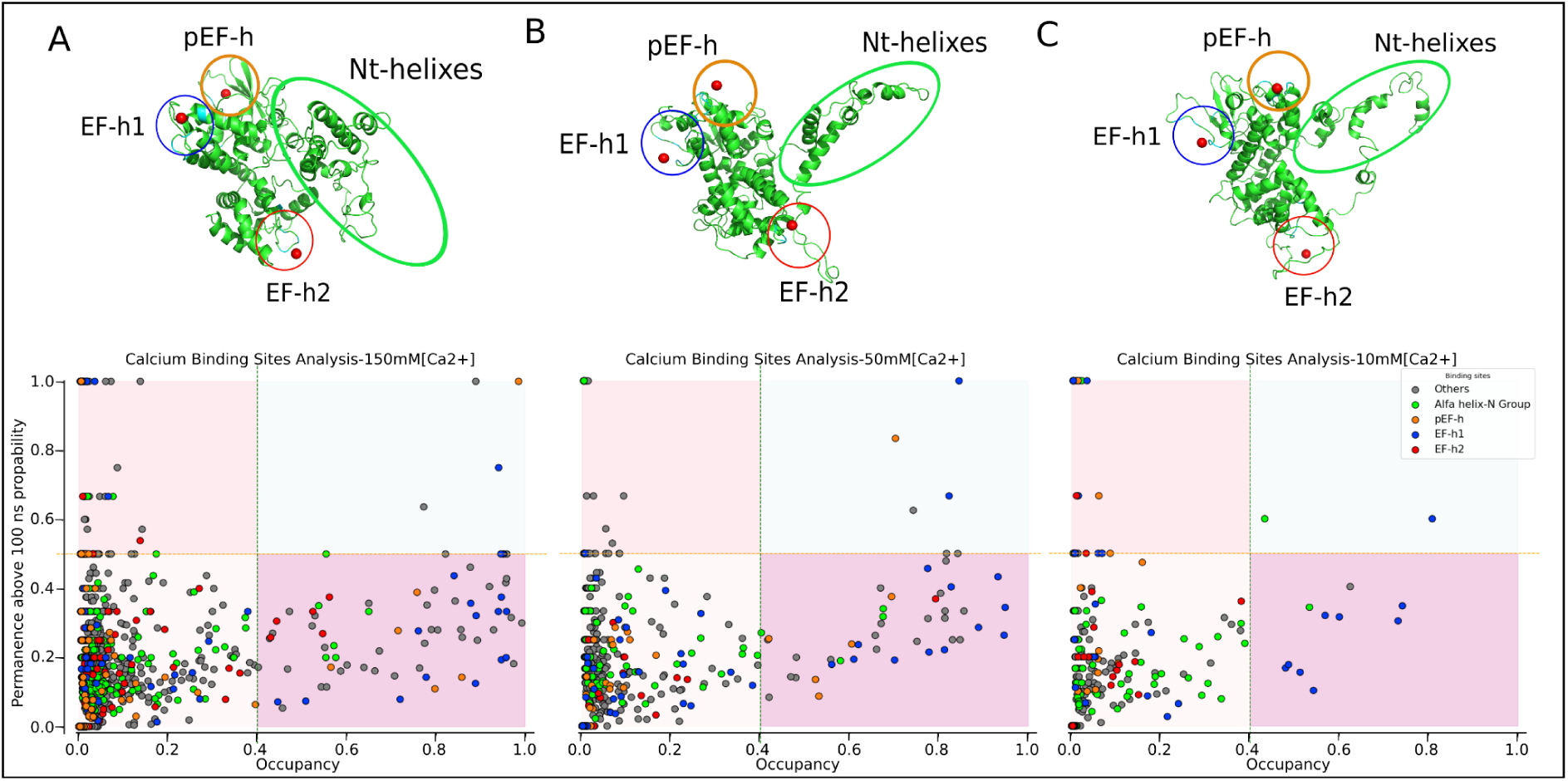
Ca^2+^-binding site analysis across different Ca^2+^ concentrations (150 mM, 50 mM, and 10 mM). Scatterplot displaying occupancy and maximum permanence time (in picoseconds) of protein residues interacting with Ca²⁺ ions. Residues are colour-coded according to their structural group: EF-h1 site (dark blue), EF-h2 site (orange), pEF-h site (orange), and N-terminal α-region (green); non-functional or low-interaction sites are shown in grey. Each panel reflects the interaction behaviour under a distinct Ca^2+^ concentration: 1A) 150 mM shows numerous residues with strong and persistent interactions, consistent with fully formed Ca^2+^-binding sites (top right quadrant); 1B) 50 mM displays moderately reduced interaction strength, with fewer residues exhibiting high occupancy and permanence, suggesting partial engagement of binding sites; 1C) 10 mM reveals generally low occupancy and short permanence times (button right quadrant), indicating the protein fails to achieve stable and complete folding under these conditions. For each concentration, a representative protein structure (fully Ca²⁺-bound) is shown, with functional regions highlighted.

The calculation of the total protein ΔG of interactions with metals across all conformers revealed that pEF-h binding to Ca^2+^ decreases the ΔG of interactions with bound metals by 6.75 ± 0.87kJ/mol (Table 2), favouring the occupancy of the remaining Ca^2+^ binding sites (EF-h1 and EF-h2) through conformational changes induced by metal binding. This decrease increases as each EF-hand is occupied (Figure 4), suggesting cooperative behaviour between the Ca^2+^ binding sites. Our results are consistent with previous proposals for EF-hand proteins, where conformational coupling enhances ion affinity and function [18,38]. Furthermore, this decrease in the protein’s ΔG of interactions with metals is accompanied by an increased proportion of exposed negatively charged residues: the percentage of negatively charged residues on the protein surface increases by 24.7%, 43.6%, and 61.6% at 150, 50, and 10 mM Ca²⁺, respectively.

**Figure 4.**
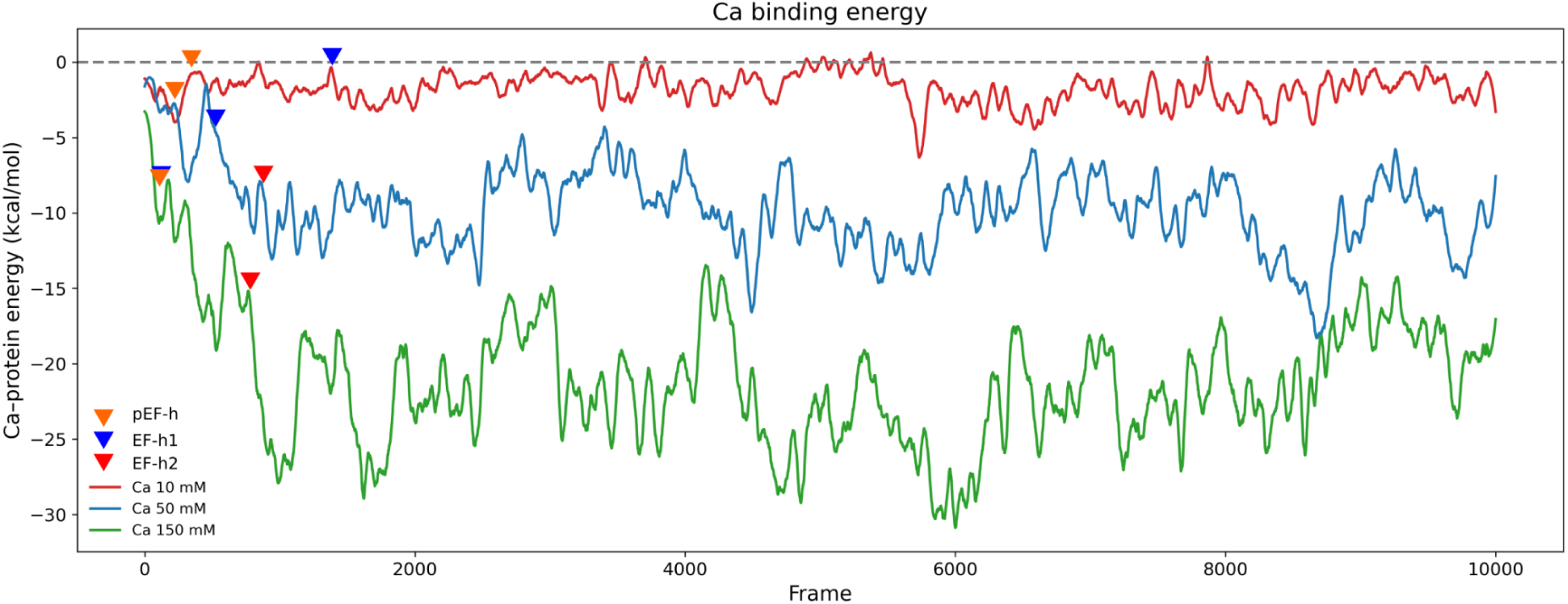
ΔG of interactions to metals per frame: in green (150 mM), blue (50 mM), and red (10 mM). Binding at pEF-h and EF-hand I and II sites reduces the ΔG of protein-metal after Ca²⁺ binding. Colorful arrows show the pEF-hand (orange), EF-hand1 (blue) and EF-hand2 (red) binding frame.

**Table 2.**
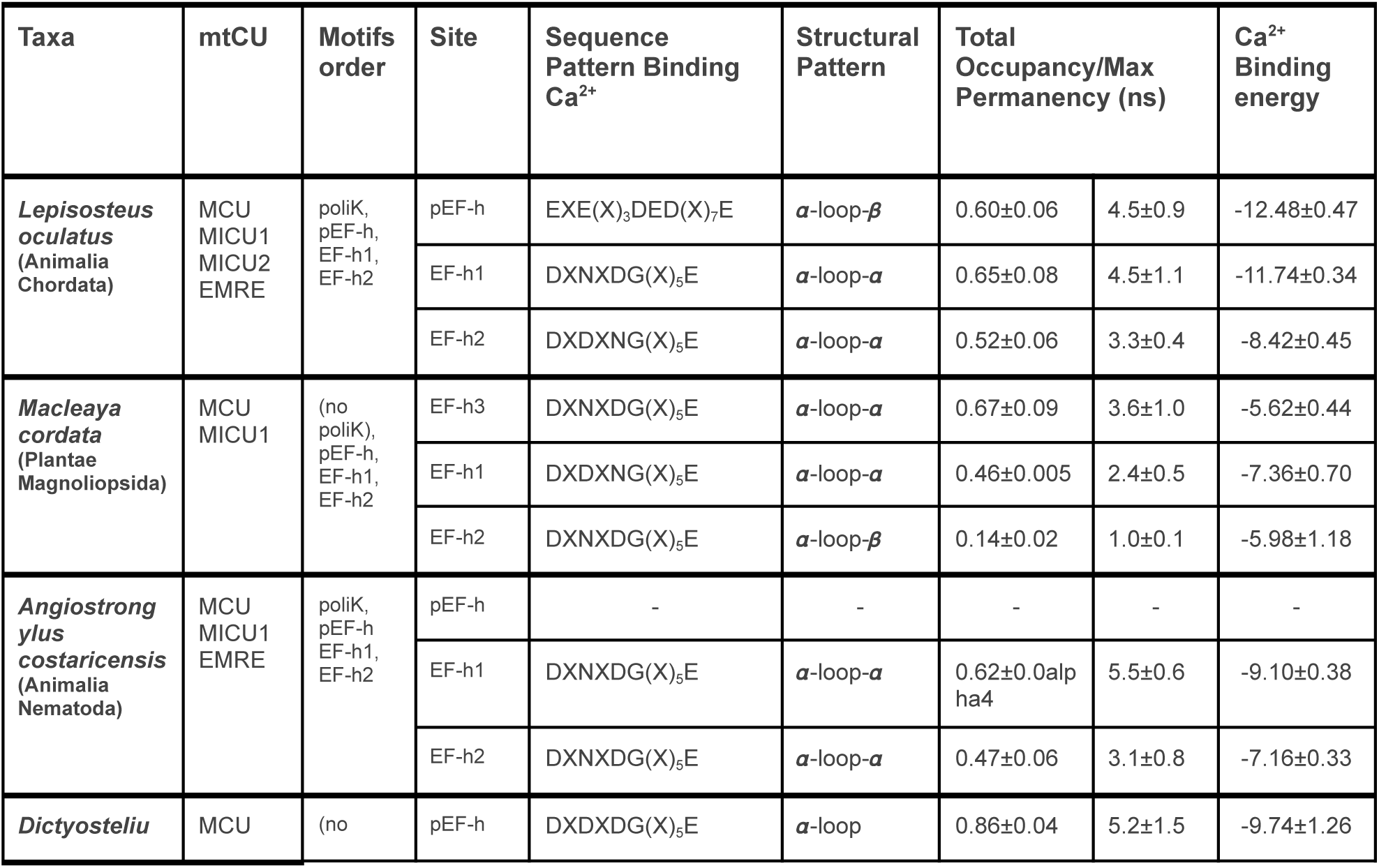
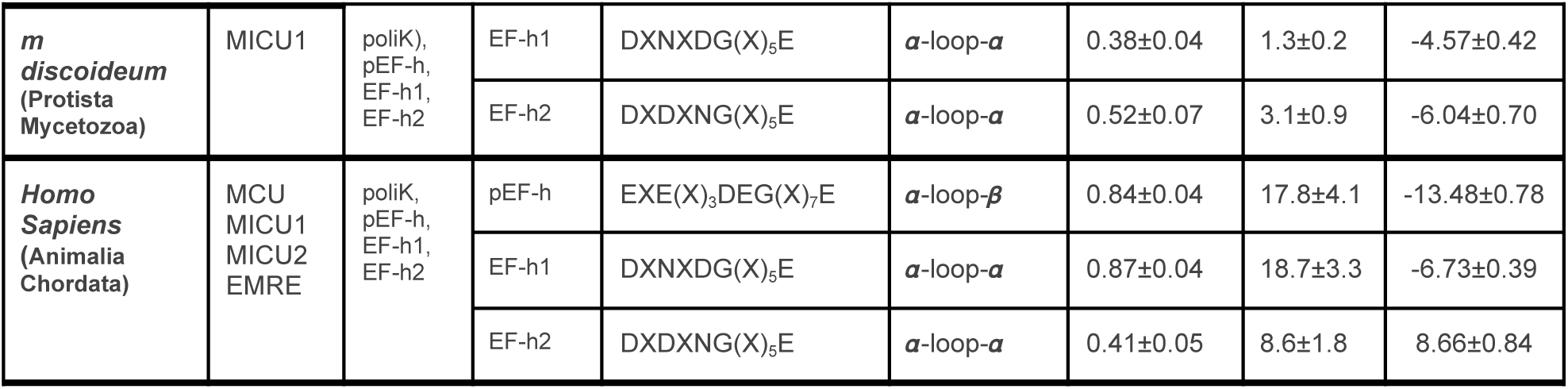
Ca²⁺-binding motifs in the pEF-h and MCU complex characteristics for sequence cluster representatives. Cluster representatives’ pEF-hand sites, including those of humans, were characterized by their sequence pattern, occupancy, permanency, and mean metal-binding energies. The mean Ca²⁺-binding energies were measured using Alphafold models from the alignments.

### In Silico Mutagenesis of MICU1 Ca²⁺-Binding Sites

To test the hierarchical order, occupancy, and permanence of Ca²⁺ binding in MICU1, we selected mutations targeting the three regulatory metal-binding regions that control channel activation. We examined how local perturbations at each site alter the temporal sequence of Ca²⁺ binding using three bibliographically annotated mutations: (i) a naturally occurring variant lacking R185 in the pEF-h site, expected to alter the earliest Ca²⁺-sensing step; (ii) the D231A single mutant and the D231A–E242K double mutant in EF-h1, designed to disrupt canonical Ca²⁺ coordination and test whether combined perturbations amplify defects in binding hierarchy and cooperativity; and (iii) the single D421H and double D421H–E432K mutants in EF-h2, which probe the role of the second EF-hand in stabilizing later stages of Ca²⁺-dependent activation. Molecular dynamics simulations of the EF-hand mutants validates what is observed experimentally. For the EF-h1 site, the single D231A mutation significantly reduced Ca²⁺ occupancy and binding permanence compared to the wild-type. However, the double mutantations D231A–E242K and D421H–E432K led to a near-complete ablation of Ca²⁺ binding specifically at the mutated EF-h site mutated (Figure 5).

**Figure 5.**
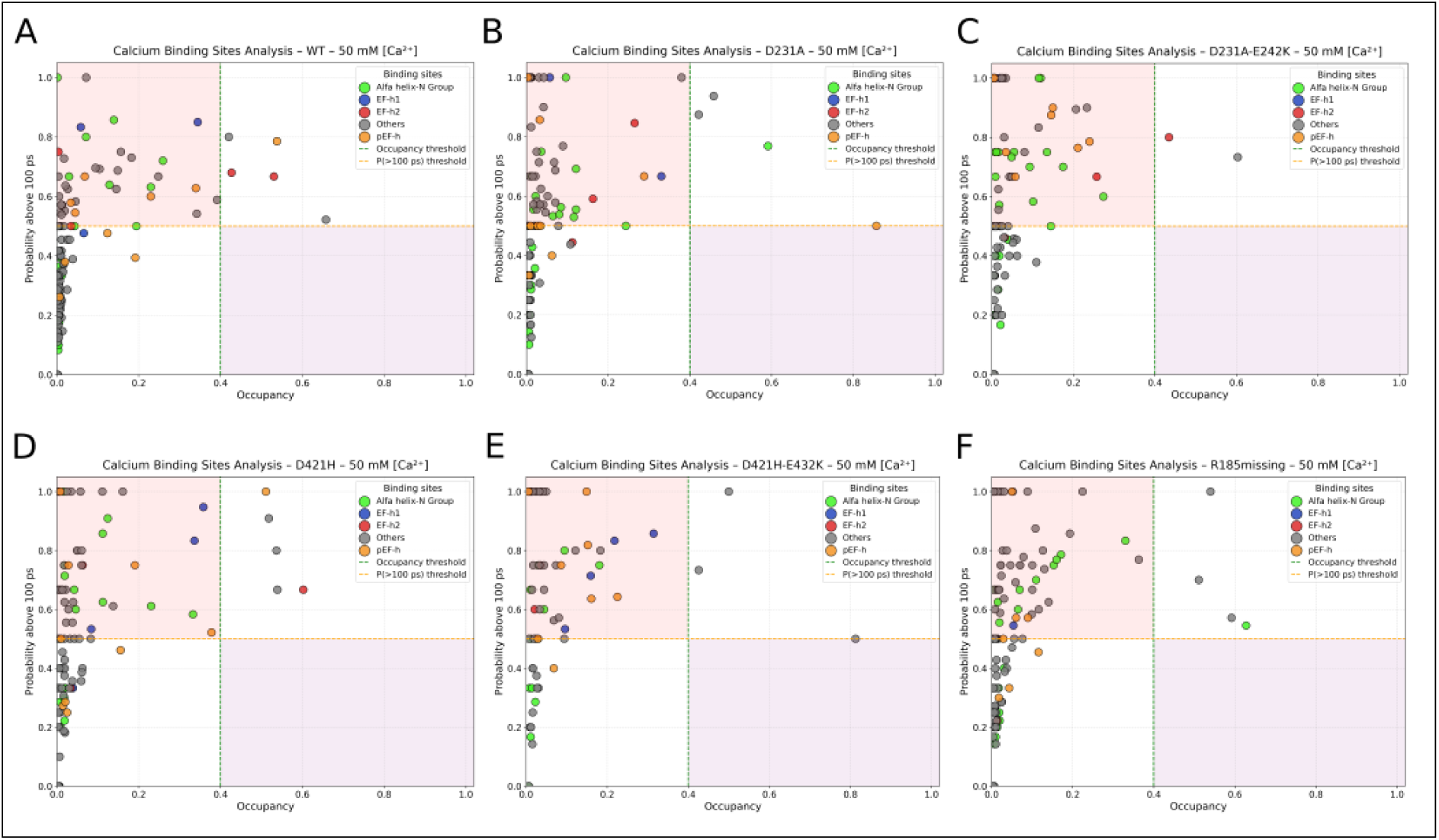
Mutations disrupt the hierarchical stability of Ca²⁺ binding. Analysis of Ca²⁺ binding metrics (occupancy vs. maximum permanence) across wild-type (A) and mutant MICU1 simulations: single mutant D231A (B), double mutant D231A–E242K (C), single mutant D421H (D), double mutant D421H–E432K (E), and the naturally occurring R185 deletion (F). Colors indicate structural groups: EF-h1 (dark blue), EF-h2 (orange), pEF-h (yellow), N-terminal α-region (green), and others (gray). The robust, high-occupancy, long-permanence interactions in wild-type (top right quadrant) are progressively eroded by single mutations (B, D). Double mutations (C, E) and the pEF-h motif deletion (F) result in near-complete loss of stable Ca²⁺ binding at the targeted sites, demonstrating the cooperative and hierarchical nature of the three-site network.

The quantitative analysis of Ca²⁺ occupancy and binding permanence in mutants compared with the wild-type form revealed a critical interdependence among the three regulatory sites, extending beyond local intra-site cooperativity. While single mutations (D231A in EF-h1 or D421H in EF-h2) primarily reduced metrics at their target site, double mutations within a canonical EF-hand pair caused propagated destabilization. Notably, the D231A-E242K double mutant in EF-h1 not only abolished Ca²⁺ binding at EF-h1 (occupancy for double D231A-E242K 0.015 ± 0.006 ns vs WT: 0.119 ± 0.152 ns) but also significantly reduced permanence at the pEF-h site (D231A-E242K 33.11 ± 23.08 ns vs WT: 157.32 ± 134.53 ns) and occupancy at EF-h2 (D231A-E242K 0.241 ± 0.202 vs WT: 0.249 ± 0.269). Similarly, the D421H-E432K double mutant in EF-h2 severely impaired its own site (occupancy for D421H-E432K 0.028 ± 0.011 ns vs WT: 0.249 ± 0.269 ns) and negatively affected binding at both EF-h1 and pEF-h. This non-local effect demonstrates that the Ca²⁺-binding sites are not independent modules in a sequence; they form an allosterically coupled network. Complete disruption of one cooperative pair perturbs the structural and energetic landscape of the entire domain, destabilizing Ca²⁺ binding at distant sites.

Consistent with its proposed role as an initial sensor, the R185 deletion variant in the pEF-h motif caused a distinct, site-specific disruption. While its effect on overall structure was minimal (comparable to wild type at non-specific sites), it caused a severe reduction in Ca²⁺ occupancy and binding permanence at its native pEF-h site (occupancy R185 deletion 0.058 ± 0.034 ns vs WT 0.159 ± 0.171 ns) and, notably, an approximately 55% decrease in occupancy at the downstream EF-h1 site (R185 deletion 0.054 ± n/a vs WT 0.119 ± 0.152). In contrast, its impact on EF-h2 was less pronounced. This pattern confirms that perturbing the early pEF-h sensor preferentially disrupts the first step in the hierarchical binding sequence, compromising the priming of subsequent sites.

This propagated failure explains the severe functional phenotypes of double mutants more comprehensively, as it reflects a collapse of the cooperative integrity required for the high-affinity, hierarchical loading of Ca²⁺ ions that drives the conformational switch.

### Evolutionary Ca²⁺-Binding Motif Conservation

To understand the contribution of Ca²⁺-binding sites to functional and evolutionary shaping, we examined the distribution of Ca²⁺-binding motifs among the 292 analyzed sequences. The global motif distribution in the alignment of the EF-h1 and EF-h2 regions revealed clear conservation patterns that further highlight their evolutionary trajectories. EF-h1 and EF-h2 are largely canonical, characterized by recurrent DXNXDG- and DLNGDGE-type motifs. These sequences retain the hallmark acidic and glycine residues typical of Ca²⁺-binding loops, which likely reflects functional specialization in maintaining robust, canonical Ca²⁺-binding (Figure 6B and 6C).

**Figure 6.**
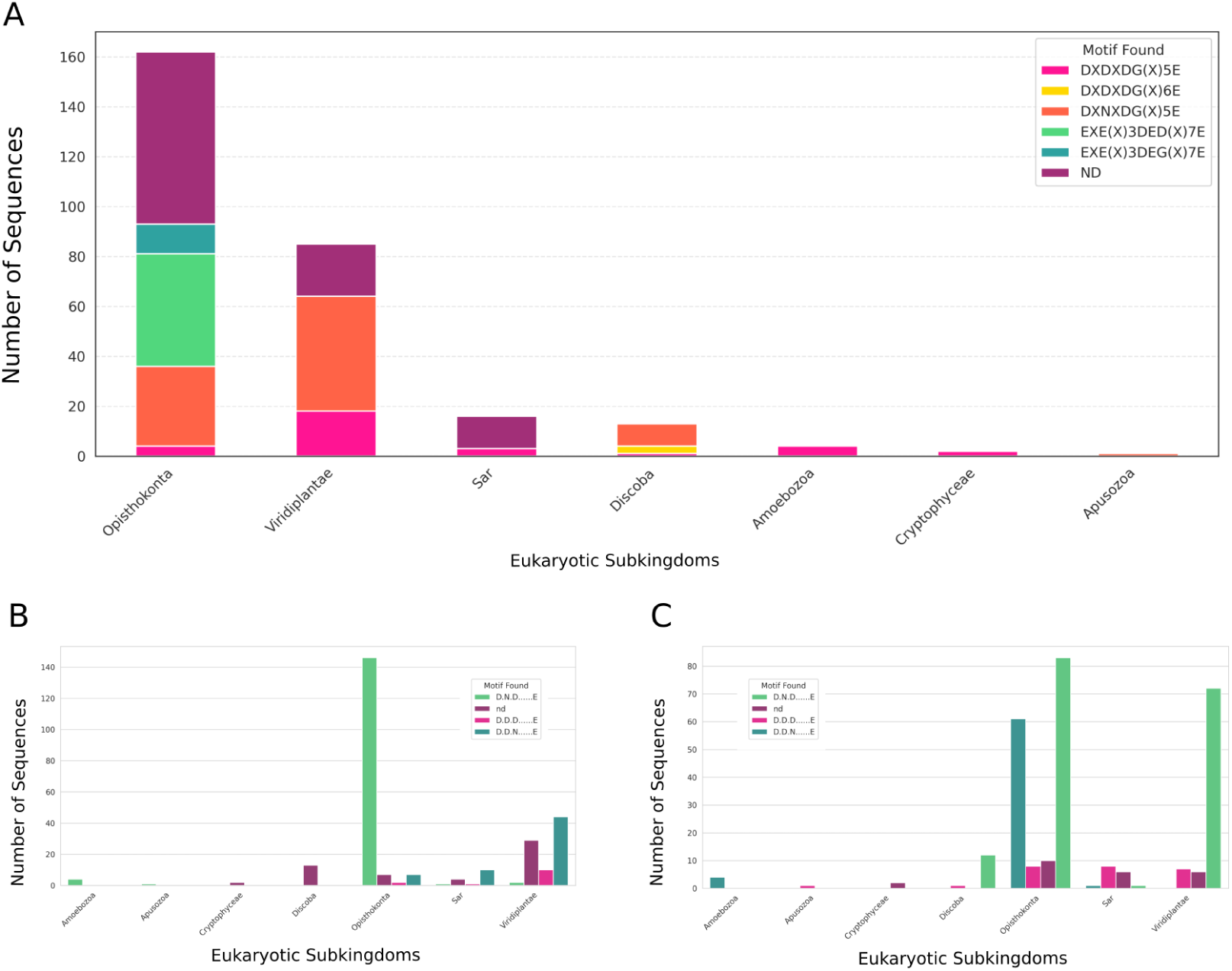
Distribution of Ca²⁺-binding motifs across major eukaryotic subkingdoms. Stacked bar plots for (A) pEF-h: bars show the number of sequences per post-Eukaryota subkingdom assigned to each conserved putative Ca²⁺-binding motif: DXNXDG(X)_₅_E (tomato), DXDXDG(X)_₅_E (deeppink), DXDXDG(X)_₆_E (gold), EXE(X)_₃_DED(X)₇E (green), and EXE(X)_₃_DEG(X)₇E (light blue). Sequences with no detectable motif are labelled nd (purple). (B) EF-h1 and (C) EF-h2: bars show the number of sequences per post-Eukaryota subkingdom assigned to each conserved putative Ca²⁺-binding motif: DXNXDG(X)_₅_E (green), DXDXDG(X)_₅_E (deeppink), and DXDXNG(X)_₅_E (light blue). Sequences with no detectable motif are labelled nd (purple).

In contrast, the pEF-h region displayed a greater variety of motifs, with increased length and larger distances between acidic and glycine residues. The EF-hand-like motif EXE(X)₃DEG(X)₇E was rare (3.77%) but notable for its presence in metazoans such as Homo sapiens. When mapped to phylogenetic groups, motif usage showed a strong lineage-specific pattern (Figure 6). The canonical DXNXDG(X)₅E motif was especially dominant in Viridiplantae and Discoba, while Opisthokonts—including animals and fungi—exhibited a broader motif distribution, encompassing all four major types. Our analysis highlights the prevalence of the DXNXDG(X)₅E motif (57.53%), which correlates with stable Ca²⁺ coordination in plants and Discoba. In contrast, the non-canonical EXE(X)₃DEG(X)₇E motif, though rare (3.77%), is exclusive to metazoans, where it supports high-affinity binding (occupancy: 0.84 ± 0.04; permanency: 17.8 ± 4.1 ns) (Table 2).

### Clustering Analysis of Ca²⁺ Binding Site Motifs

To investigate the relationship between motif composition and site functionality, we performed a clustering analysis on the 292 motifs identified in the alignment and its structural information (Figure 7). This revealed strong phylogenetic segregation: clusters 0 and 2 were predominantly composed of the canonical DXNXDG(X)₅E and DXDXDG(X)₅E motifs, prevalent in Viridiplantae and Discoba, whereas cluster 3 was highly enriched for the non-canonical EXE(X)₃DEG(X)₇E and related EXE(X)₃DED(X)₇E motifs characteristic of Opisthokonta. This partitioning correlated with functional divergence: the DXN/DXD-dominated clusters showed weaker Ca²⁺-binding energies, consistent with a role as dynamic sensors, while the EXE-enriched cluster exhibited the strongest binding affinities, reflecting specialization as high-fidelity Ca²⁺ gatekeepers in metazoans. Moreover, the prevalence of the poliK domain was highest in the basal EXE-motif clusters but substantially reduced in the derived DXD-motif cluster, suggesting that the ancestral poliK-associated regulatory mechanism was often modulated or refined in lineages that evolved the high-affinity EXE motif and the EMRE subunit. Together, these results support an evolutionary model in which pEF-hands evolved from simpler, dynamic sensors in early eukaryotes to sophisticated, high-affinity Ca²⁺ gatekeepers in complex multicellular organisms.

**Figure 7.**
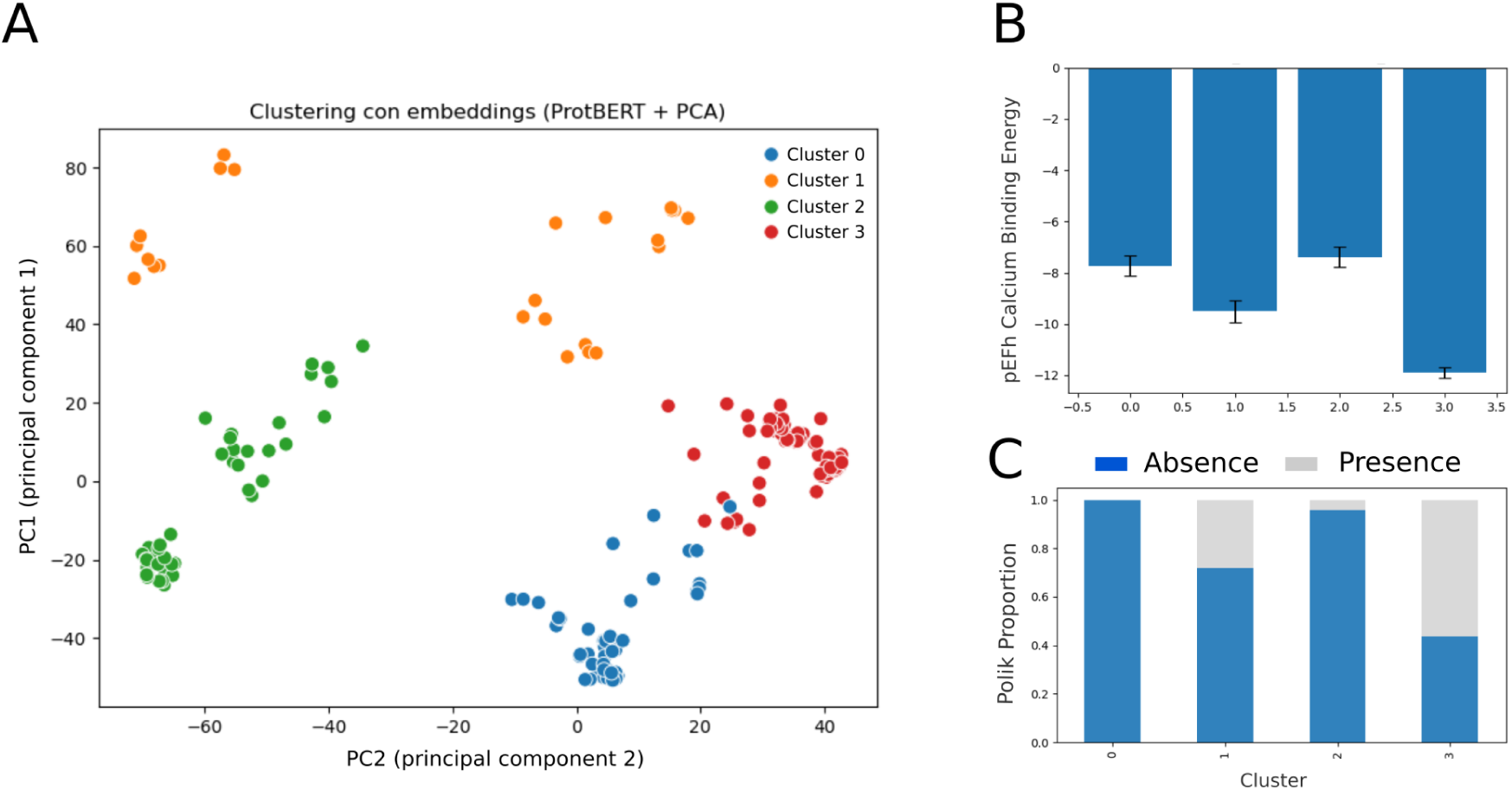
Clustering analysis of pEF-hand motifs across eukaryotes. A) Sequence-based clustering of 292 pEF-hand motifs identifies four distinct clusters (0–3), coloured as indicated. Cluster 0 (blue) is dominated by non-canonical DXN/DXD motifs (D.N.D……E and D.D.D E), prevalent in Viridiplantae and Discoba. Cluster 1 (orange) contains mostly DXD motifs with rare EXE-like motifs, representing a potential evolutionary transition. Cluster 2 (green) is primarily DXN/DXD but includes a small fraction of canonical EXE(X)_₃_DEG(X)_₇_E motifs, suggesting the emergence of canonical motifs in basal lineages. Cluster 3 (red) is enriched for canonical EXE motifs, characteristic of Opisthokonta and metazoans. B) Predicted Ca²⁺-binding energies for each cluster. Cluster 3 exhibits the strongest binding (lowest energies), consistent with functional specialization. C) Proportion of sequences containing the PoliK domain in each cluster. PoliK prevalence is highest in the basal DXN/DXD-rich clusters (0 and 2), reduced in the transitional Cluster 1, and lowest in the canonical EXE-enriched Cluster 3, suggesting an evolutionary shift in regulatory mechanisms.

In EF-h1 and EF-h2, the diversity of Ca²⁺-binding motifs was lower, mostly showing canonical motifs with similar energies within clusters, but lower compared to pEF-hand motifs.

### Structural and Evolutionary Constraints Over Ca²⁺-Binding Sites in Biology Systems

To investigate variations in MICU1 Ca²⁺-binding sites and potential functional adaptations across species, we constructed a phylogenetic tree and performed sequence clustering to select representative proteins for detailed structural and functional comparison of their pEF-hand and EF-hand motifs. Each representative, including human MICU1, was characterized using molecular dynamics simulations at 150 mM Ca²⁺ to assess Ca²⁺-binding affinities and reveal lineage-specific adaptations in EF-hand coordination. Our analysis indicates that the pEF-hand region (residues 184–200), essential for Ca²⁺ coordination, exhibits significant evolutionary divergence across eukaryotes (Table 2).

In Amoebozoa, this region retains a canonical EF-hand motif with the classical helix–loop–helix structure, whereas in mammals, despite broader structural divergence across Eukarya, the pEF-hand sequence remains highly conserved, underscoring its critical role in Ca²⁺ regulation [37]. In contrast, plants such as Macleaya cordata maintain the overall α–loop–α fold but show substantial sequence variation, resulting in the loss of canonical EF-hand residues and reduced Ca²⁺-binding occupancy (0.14 ± 0.02) compared to metazoans, suggesting potential functional adaptations or alternative regulatory roles, consistent with observations in other systems [11,47,48].

The DxDxDG motif, traditionally associated with EF-hand structures, is now identified in various structural contexts, indicating evolutionary repurposing rather than a single ancestral origin [39,47]. The EF-hand function is primarily structural in bacteria and regulatory in eukaryotes [35,48]. However, exceptions have been observed in some eukaryotes, where the EF-hand occasionally displays high affinity, resulting in a purely structural role [44]. Our comparative analysis reveals notable lineage-specific differences in Ca²⁺-binding dynamics. In humans, the pEF-hand belongs to Cluster 3, characterized by canonical EXE motifs, and exhibits optimised high-affinity binding (−13.48 ± 0.78 kcal/mol), consistent with its role as a precise regulator of mitochondrial Ca²⁺ uptake. In contrast, in plants such as Macleaya cordata, pEF-hands are predominantly DXN/DXD motifs corresponding to Clusters 0 and 2, retaining the α-loop-α fold but showing weaker Ca²⁺-binding, with low occupancy (0.14 ± 0.02), short permanency (1.0 ± 0.1 ns), and less negative binding energies (-5.62 ± 0.44 kcal/mol). This suggests either a secondary Ca²⁺-sensing function or a divergent regulatory mechanism.

In Amoebozoa, exemplified by Dictyostelium discoideum, pEF-hands are predominantly found in Cluster 0, showing moderate Ca²⁺ permanency (5.2 ± 1.5 ns) despite high occupancy (0.86 ± 0.04) and intermediate binding energy (-9.74 ± 1.26 kcal/mol), which suggests dynamic Ca²⁺-sensing functions. The absence of EMRE in this lineage is associated with less energetically favourable interactions compared to vertebrates. In nematodes such as Angiostrongylus costaricensis, EF-hand motifs display intermediate occupancy (0.47–0.62) and permanency (3.1–5.5 ns) with moderately strong binding energies (-9.10 ± 0.38 kcal/ -7.16 ± 0.33 kcal/mol), reflecting functional divergence in Ca²⁺ regulation [36–38].

Taken together, these observations support an evolutionary model in which pEF-hands evolved from dynamic Ca²⁺ sensors in basal eukaryotes, characterized by DXN/DXD motifs (Clusters 0 and 2), towards high-affinity canonical EXE motifs (Cluster 3) in metazoans. The lower binding energies and shorter permanency observed in plants and Amoebozoa indicate more transient Ca²⁺ interactions, whereas vertebrates display stable, high-affinity coordination suitable for precise regulation of mitochondrial Ca²⁺ uptake. Overall, these results underscore the interplay between motif sequence, structural divergence, and co-evolution with regulatory subunits such as EMRE in shaping Ca²⁺-binding dynamics across eukaryotic lineages.

## Conclusion

Our study provides new structural and evolutionary insights into the role of MICU1 in regulating the mitochondrial Ca²⁺ uniporter complex, deepening our understanding of its function in Ca²⁺ signaling and energy homeostasis. Through sequence analysis, structural modeling, and molecular dynamics simulations, we characterized Ca²⁺-induced conformational changes in MICU1, revealing substantial shifts in global structure, charge distribution, and cavity architecture upon Ca²⁺-binding.

One key finding was the cooperative behavior among Ca²⁺-binding sites, revealed by the progressive decrease in ΔG of protein–metal interactions with sequential Ca²⁺-binding. We observed that initial binding at the pEF-hand (pEF-h) lowered ΔG by approximately 6.2 kJ/mol, which facilitated subsequent occupancy of EF-hand I and II through conformational coupling. This cooperative effect was further reinforced by increased exposure of negatively charged surface residues, which expanded with rising Ca²⁺ concentration. Together, these observations highlight how Ca²⁺ coordination remodels MICU1’s electrostatic surface to enhance ion binding and stabilize its active conformations.

Furthermore, we identified and characterized the pEF-hand (pEF-h) motif, spanning residues 184–200. Unlike canonical EF-hand motifs, the pEF-h adopts a helix–loop–β-sheet fold while maintaining functional Ca²⁺-binding capability. Molecular dynamics simulations indicate that this motif acts as a dynamic Ca²⁺ sensor, triggering structural transitions that enable subsequent binding at EF-hand I and II, thus playing a central role in MICU1’s on–off regulatory mechanism. However, future experimental validation, using fluorescence assays, spectroscopy, or targeted mutagenesis, will be essential to complement our computational predictions, test cooperativity of the binding motifs and observe the full folding–unfolding transitions described in the present work.

Importantly, our in silico mutagenesis study provides a direct mechanistic link between this cooperative model and its functional consequences. We elucidate the molecular mechanism underlying the experimental observations, showing that the three Ca²⁺-binding sites form an allosterically coupled network. Disruption of one site propagates to destabilise the others. Specifically, double mutations within a canonical EF-hand (D231A-E242K in EF-h1 or D421H-E432K in EF-h2) not only abolish local Ca²⁺ binding but also significantly reduce occupancy and permanence at the distal, non-mutated sites. This quantifies the global cooperativity of the system, explaining why single mutations often yield mild phenotypes, while combined mutations produce severe, synergistic loss of function. The collapse of the entire network upon destabilisation of one node underscores how MICU1 integrates multiple Ca²⁺-binding events into a unified, high-fidelity on-off switch, a design principle that balances sensitivity with robustness in mitochondrial Ca²⁺ signalling.

Evolutionary analyses of 292 MICU1 sequences revealed strong phylogenetic segregation of Ca²⁺-binding motifs. The canonical DXN/DXD-type motifs of EF-h1 and EF-h2 are highly conserved across eukaryotes, reflecting strong functional constraints. In contrast, the pEF-h region exhibits greater motif diversity, with DXN/DXD variants predominating in basal lineages such as Viridiplantae and Discoba, and non-canonical EXE(X)₃DEG(X)₇E motifs emerging exclusively in metazoans (Opisthokonta). This motif specialization suggests evolutionary tuning of Ca²⁺-binding affinity: the dynamic pEF-h acts as an early sensor, while the metazoan-specific EXE motif in the same region may enable tighter, more stable Ca²⁺ coordination. This combination supports a refined activation model, in which initial sensing by the pEF-h is followed by high-affinity locking via the EXE motif, aligning with the more complex Ca²⁺ signaling demands of multicellular organisms.

The appearance of the EMRE subunit coincides with this transition, suggesting the co-evolution of structural motifs and regulatory subunits to achieve tighter control of mitochondrial Ca²⁺ uptake. In taxa lacking EMRE, pEF-h interactions are weaker or transient, implying reduced regulatory precision or alternative Ca²⁺-sensing mechanisms.

Overall, our findings show that MICU1’s Ca²⁺-binding dynamics are shaped by the interplay of motif sequence, structural divergence, and co-evolution with regulatory subunits. The pEF-h is a key evolutionary innovation that initiates early Ca²⁺-sensing and coordinates the activation of the canonical EF-h1 and EF-h2 sites. Importantly, the emergence of lineage-specific motifs, such as EXE in metazoans, could reflect evolutionary adaptations to distinct physiological requirements. By integrating evolutionary analysis with structural modeling and molecular dynamics, we reveal how MICU1 alone can function as a Ca²⁺-dependent molecular switch, providing a mechanistic framework for its threshold-setting and cooperative behavior, and a blueprint for understanding similar regulatory mechanisms in other Ca²⁺-sensing proteins.

## Materials and Methods

### Structure Modelling and Molecular Dynamics

To investigate the conformational dynamics of MICU1, we performed structural modelling using AlphaFold2 [25] (total pDDLT > 70), based on the incomplete three-dimensional structure available in the Protein Data Bank (PDB ID: 4NSC). The model was generated using default AlphaFold2 parameters.

Molecular dynamics simulations were carried out using GROMACS [1], to explore structural changes in human MICU1 at varying Ca²⁺ concentrations (0, 10, 50, and 150 mM). The selected concentrations assume that the close proximity between MICU and the Ryanodine Receptor generates a high Ca²⁺ microenvironment [12,45], and thus the concentrations required to observe the conformational changes supporting protein function will be higher than those in the cytosol.

Each simulation was run for 100 nanoseconds, using an all-atom representation with the CHARMM36 force field and the TIP3P water model [23]. The v-rescale thermostat was used to maintain the temperature (T = 310 K) with a relaxation time of 0.1 ps, and Berendsen pressure coupling (1 atm) with a relaxation time of 1 ps was applied in the NPT ensemble [4]. Pressure was coupled isotropically in the XY plane and the Z direction. The time step was 2 fs. The last frame of this equilibrated system was used as the initial configuration for a second simulation. The Particle Mesh Ewald (PME) method was used to treat electrostatic interactions [14,20], with a real space cut-off of 4 nm, a grid spacing of 0.16 nm, and cubic interpolation. We observed that 100 ns simulations are enough for capturing the Ca^2+^ binding allosteric cycle.

The starting conformation for the fully unbound form was taken from the structure at 0 mM Ca²⁺, while the fully Ca²⁺-bound form was defined as the structure with occupied EF-h sites. For each cluster representative protein in the phylogenetic tree obtained from the evolutionary analysis, we structurally characterized them through 30 nanoseconds molecular dynamics simulations (at 150 mM Ca²⁺), and calculated the metal-binding affinities using FoldX across all simulation frames [50].

### Structural Analysis

Janus analysis was performed according to the methodology described by Madhurima et al. [32]. he structural characterisation of MICU1 in both bound and unbound cavity forms was carried out using CaviDB [54] to analyse cavity properties, including volume, hydrophobicity, and solvent accessibility. Key structural parameters, such as root mean square deviation (RMSD), accessible surface area (ASA), and residue–residue contacts, were calculated to assess conformational changes upon Ca²⁺-binding. We also analysed the sequence composition and charge distribution with localCIDER [22].

For comparative analysis, unbound and fully Ca²⁺-bound conformations were structurally aligned using Chimera [43], enabling quantification of residue distance changes.

The occupancies and permanency of Ca²⁺ ions within the MICU1 pEF-hand and EF-hand motifs were computed from molecular dynamics simulations. total occupancy was defined as the fraction of simulation time during which a Ca²⁺ ion remained coordinated within the canonical EF-hand motif binding site, averaged across replicates. Permanency was calculated as the maximum continuous time (in nanoseconds) that a Ca²⁺ ion remained bound during the trajectory.

The metal binding energy was calculated using FoldX [50] by applying the MetalScan command to all frames of the trajectory. FoldX estimates the binding energy of Ca²⁺ ions to protein motifs using a semi-empirical energy function that includes van der Waals interactions, solvation, and metal coordination terms. The minimum binding energy (kcal/mol) was reported as the lowest energy value observed during the entire simulation.

All structural and dynamic data were processed using custom Python scripts, incorporating the MDAnalysis, Seaborn, Pandas, and SciPy libraries [34,55,57].

Hinge residues were identified by comparing backbone dihedral angles (φ, ψ) between the unbound and Ca²⁺-fully bound MICU1 structures after superposition [9,46]. Dihedral differences were computed with angular periodicity correction, and the total dihedral change per residue was defined as the Euclidean combination of Δφ and Δψ. Residues with total dihedral variation above the 75th percentile of the global distribution were classified as hinge residues. Regional enrichment of hinge residues and backbone rearrangements was evaluated for functional domains using Welch’s t-tests and effect size estimates (Cohen’s d). Solvent accessible surface area (ASA) was calculated for each residue of MICU1 in both unbound and Ca²⁺-fully bound conformations using the Shrake–Rupley algorithm implemented in BioPython [52]. For each functional region, the median and standard deviation of residue ASA values were computed for the unbound and full bound conformations. The change in ASA (ΔASA) upon Ca²⁺ binding was defined as the difference between the median ASA in the bound and unbound conformations. Regions exhibiting positive ΔASA indicate increased solvent exposure upon Ca²⁺ binding, whereas negative ΔASA reflects conformational collapse or burial of residues.

### In Silico Mutagenesis Analysis

Previous experimental studies have shown that single mutations in EF-h1 often produce mild or undetectable functional effects, while combined mutations generate robust phenotypes, highlighting the cooperative nature of Ca²⁺ sensing in MICU1. In particular, double mutants in EF-h1 impair mitochondrial Ca²⁺ uptake more efficiently than isolated substitutions, supporting the idea that disruption of a single coordinating residue may be partially compensated by the remaining network of interactions. Based on these observations, we examined: (i) a naturally occurring variant lacking R185 in the pEF-h site, which is expected to alter the earliest Ca²⁺-sensing step [27]; (ii) the D231A single mutant and the D231A–E242K double mutant in EF-h1, designed to disrupt canonical Ca²⁺ coordination and test whether combined perturbations amplify defects in binding hierarchy and cooperativity [26,42]; and (iii) the single D421H and double D421H-E432K mutant in EF-h2, which probes the role of the second EF-hand in stabilizing later stages of Ca²⁺-dependent activation [26,33,42].

By analyzing these variants, we aimed to determine how local perturbations at each site alter the temporal sequence of Ca²⁺ binding, the stability of ion occupancy, and the permanence of metal–protein interactions, thereby providing a mechanistic framework to understand how MICU1 integrates multiple Ca²⁺-binding events into a cooperative on–off regulatory switch for the mitochondrial Ca²⁺ uniporter. Thus, we modeled all the variants from the wild-type unbound structure using AlphaFold. For all mutants, we ran simulations of 30 nanoseconds using an all-atom representation with the CHARMM36 force field and the TIP3P water model. We also performed structural analyses of each functional domain and directly compared with the wild type in both unbound and Ca²⁺-bound conformations, following the same methodological framework used for the wild type.

### Evolutionary Analysis

A total of 292 sequences were extracted from the OMA database [3], excluding sequences with substantial deletions or truncations in the MICU1 functional domains, and aligned using Clustal [28] (with the BLOSUM62 scoring matrix, gap opening penalty of 10, gap extension penalty of 0.1, end gap separation penalty and transition weighting of 0.5) before sequence clustering with CD-HIT [24], using 90% identity, and a window of 4, and 90% coverage, as parameters. This grouped sequences by identity to track the sequential and structural conservation of the pEF-h motif.

To investigate the evolutionary conservation of MICU1, we performed phylogenetic inference using IQ-TREE with maximum likelihood methods [36]. Model selection was performed automatically by IQ-TREE using ModelFinder, which identified JTT+R5 as the best-fit substitution model. To assess functional conservation, we calculated the metal binding energy for the pEF-h and EF-hand sites in the homologues using FoldX’s metal binding command [50], over the entire trajectory from Ca^2+^-free to fully bound. We used the minimum metal binding energy as a reference for interspecies comparison of the calculated energies. The distribution of motif sequence patterns was analyzed using custom Python scripts, incorporating the Seaborn, Pandas, and Numpy libraries [34,55,57].

### Clustering Analysis of pEF-hand Motifs

To identify evolutionary patterns among the 292 pEF-hand sequences, we performed comprehensive clustering analysis that integrated both numerical features and sequence-based characteristics. For each sequence, we extracted quantitative descriptors, including Ca²⁺-binding energies, occupancy metrics, and structural parameters. We also extracted simple sequence features, such as motif lengths and position-specific amino acid compositions. Additionally, sequence features were additionally encoded using deep learning-based embeddings generated with ProtBERT (Rostlab/prot_bert) to capture evolutionary and structural relationships, resulting in 1024-dimensional vector representations for each motif region. After imputing missing values using mean substitution and standardisation, we applied K-means clustering with k = 4, determined through empirical evaluation. The resulting clusters were visualized using principal component analysis (PCA) and validated by examining the distribution of key biological features across clusters, including the presence of Polik domains and other mtCU component binding sites. All clustering analyses used fixed random seeds (random_state=42) and 10 K-means initializations (n_init=10) to ensure reproducibility. This multi-faceted approach ensured that clusters reflected both functional properties and evolutionary relationships within the pEF-hand repertoire. The metal binding energy for the pEF-h and EF-hand sites in the homologues was measured using FoldX’s metal binding command [50], and distributions were analysed using custom Python scripts incorporating the Seaborn, Pandas, and Numpy libraries [34,55,57].

### Statistical analysis

All statistical analyses were performed using Python library *scipy*. All data were first tested for normality with the D’Agostino and Pearson’s normality test. Homoscedasticity was tested using Levene’s test. T-test or Mann–Whitney test were carried out for pairwise comparisons. For multigroup comparisons, one-way ANOVA or the non-parametric equivalent Kruskal–Wallis test was used when appropriate. Tukey’s test and Dunn’s multiple comparisons test were used when appropriate.

## Authors’ contributions

LMS and AJVR carried out the bioinformatic analysis. TG carried out the evolutionary analysis conceptualization. NP carried out the linear motifs analysis conceptualization. AJVR and LMS conceived of the study, and participated in its design and coordination. GP and MSF carried out the project administration and funding acquisition. All authors participated in drafting the manuscript, as well as the final manuscript reading and approval.

## Acknowledgements

LMS, AJVR, NP, GP and MSF are researchers from Consejo Nacional de Investigaciones Científicas y Técnicas (CONICET). This work was supported by Universidad Nacional de Quilmes (PUNQ 918/22). This work used computational resources from UNC Supercómputo (CCAD) – Universidad Nacional de Córdoba (https://supercomputo.unc.edu.ar), which are part of SNCAD, República Argentina.

## Data availability

All data can be accessed at the CONICET open data repository: https://datosdeinvestigacion.conicet.gov.ar/handle/11336/278960. Due to space limitations, the uploaded trajectories are samples representing the conformational changes, selected using normalized RMSD distribution for frame selection. Additional data will be shared upon request.

